# Drug REpurposing using AI/ML tools - for Rare Diseases (DREAM-RD): A case study with Fragile X Syndrome (FXS)

**DOI:** 10.1101/2020.09.25.311142

**Authors:** Kavitha Agastheeswaramoorthy, Aarti Sevilimedu

## Abstract

Drug repositioning is emerging as an increasingly relevant option for rare disease therapy and management. Various methods for identifying suitable drug candidates have been tried and range from clinical symptomatic repurposing to data driven strategies which are based on the disease-specific gene or protein expression, modification, signalling and physiological perturbation profiles. The use of Artificial Intelligence (AI) and machine learning algorithms (ML) allows one to combine diverse data sets, and extract disease-specific data profiles which may not be intuitive or apparent from a subset of data. In this case study with Fragile X syndrome and autism, we have used multiple computational methodologies to extract profiles, which are then combined to arrive at a comprehensive signature (disease DEG). This DEG was then used to interrogate the large collection of drug-induced perturbation profiles present in public databases, to find appropriate small molecules to reverse or mimic the disease-profiles. We have labelled this pipeline **D**rug **Re**purposing using **A**I/**M**L tools - for **R**are **D**iseases (DREAM-RD). We have shortlisted over 100 FDA approved drugs using the aforementioned pipeline, which may potentially be useful to ameliorate autistic phenotypes associated with FXS.

## Introduction

Fragile X syndrome is a rare disease that occurs due to a loss of function of the Fragile-X mental retardation protein (FMRP) in humans (1,2) Symptoms characteristic of FXS include craniofacial elongation, macroorchidism, elongated limbs, cognitive deficits and behavioural abnormalities like anxiety, irritability and aggression (3). FXS is the leading monogenic cause of Autism spectrum disorders (ASD) (4).

FMRP is known to be a selective RNA binding protein, and contains two independent RNA binding domains. In adult neurons, FMRP is known to interact with several mRNAs (5,6) as well as associate with elongating polyribosomes (7,8). FMRP is believed to regulate the translation of these mRNAs by various mechanisms, and influences their transport to dendritic shafts, spines and growth cones thus directly affecting synaptic plasticity and development (9). FMRP has been shown to associate with over 800 mRNAs in adult neurons, and loss of FMRP is therefore likely to affect the functional network of a large number of gene products and signalling cascades.

The disruption of a key protein (FMRP) that drives an important function (synaptic development and plasticity), and directly impacts hundreds of gene products (via translation elongation), is likely to result in measurable and profound changes at the level of gene expression. These changes will include those that are a direct result of the loss of FMRP function, and compensatory changes to adjust for the said loss. And these changes form the primary starting point, of a cascade that results in cellular phenotypes which then manifest as behavioural symptoms in an FXS patient. Therefore the differential gene expression patterns present in FXS patients as compared to the normal populace can be used to identify potential intervention points for therapy.

Rare genetic diseases like FXS, are aetiologically well defined, but therapeutically challenging because of the nature and complex cellular roles of FMRP. The diverse array of symptoms stem from disparate pathways that are dysregulated, therefore the traditional target based drug discovery approaches have not been successful (10,11). In order to tackle such a problem a) multiple targets or pathways that drive the core “phenotypes” of FXS need to be identified and b) therapeutics (or combinations) which impact multiple targets need to be prioritised.

Drug repurposing has been the favoured approach to discover therapies for rare diseases, because of the significant time and cost saving it offers. Repurposing involves the use of a drug with proven efficacy for a particular indication and regulatory approval, for a different indication, irrespective of the original target. In order to repurpose a drug for a new indication, the consequence of the drug or the disease on the entire cellular network (transcriptome, proteome, signalling network or all of these) must be known, in order to extract potentially new overlaps or non-obvious connections, which may not have been considered important for each (drug or new indication) in isolation. The larger the number of data features and sample numbers available for generating such a comprehensive map, the stronger the likelihood of identifying a drug repurposing candidate(12,13). However, gene expression profiles are the only type of data available for most diseases and drug treatments and therefore form the basis of a large part of repurposing efforts in current times.

The general workflow of computational methods for drug repurposing, involves the extraction of a disease specific gene expression signature from patient transcriptomics data using computational methods. The signature becomes the input parameter set to filter and find drug induced perturbations that are similar (mimic) or opposite in direction (reverse) and to find dysregulated pathways which may reveal non-obvious drug targets. The drug specific gene expression profiles are inferred from cell line data available from public datasets like CMAP (14) and LINCS (15), or experimental datasets deposited in GEO (16).

Several strategies for drug repurposing have been tried in the case of FXS, such as clinical phenotype matching, based on molecular mechanisms and empirical observations and a number of these are summarised in Tranfaglia et al (17). Two key points differentiate our approach from the previously reported studies for FXS repurposing. The first, is the use of gene expression profiles from FXS as well as autism patients, in order to derive a disease signature that represents the autistic spectrum of phenotypes specific to FXS. Since FMRP may have broad roles during early development and translation regulation, and the aetiology of ASD could be varied, combining the two may add weight to and highlight the truly causative changes underlying the autistic phenotypes. Second, we have combined ML methods to the traditional statistical methods of gene expression analysis, in order to derive a robust and comprehensive disease signature of differentially expressed genes (DEGs) from relatively small amount of data. ML algorithms are increasingly favoured for their computational efficiency in extracting important features from high dimensional data.

## Methods

### Datasets

Gene expression data was obtained from NCBI GEO and the accession number are listed in Table 1.

**Table 1.**
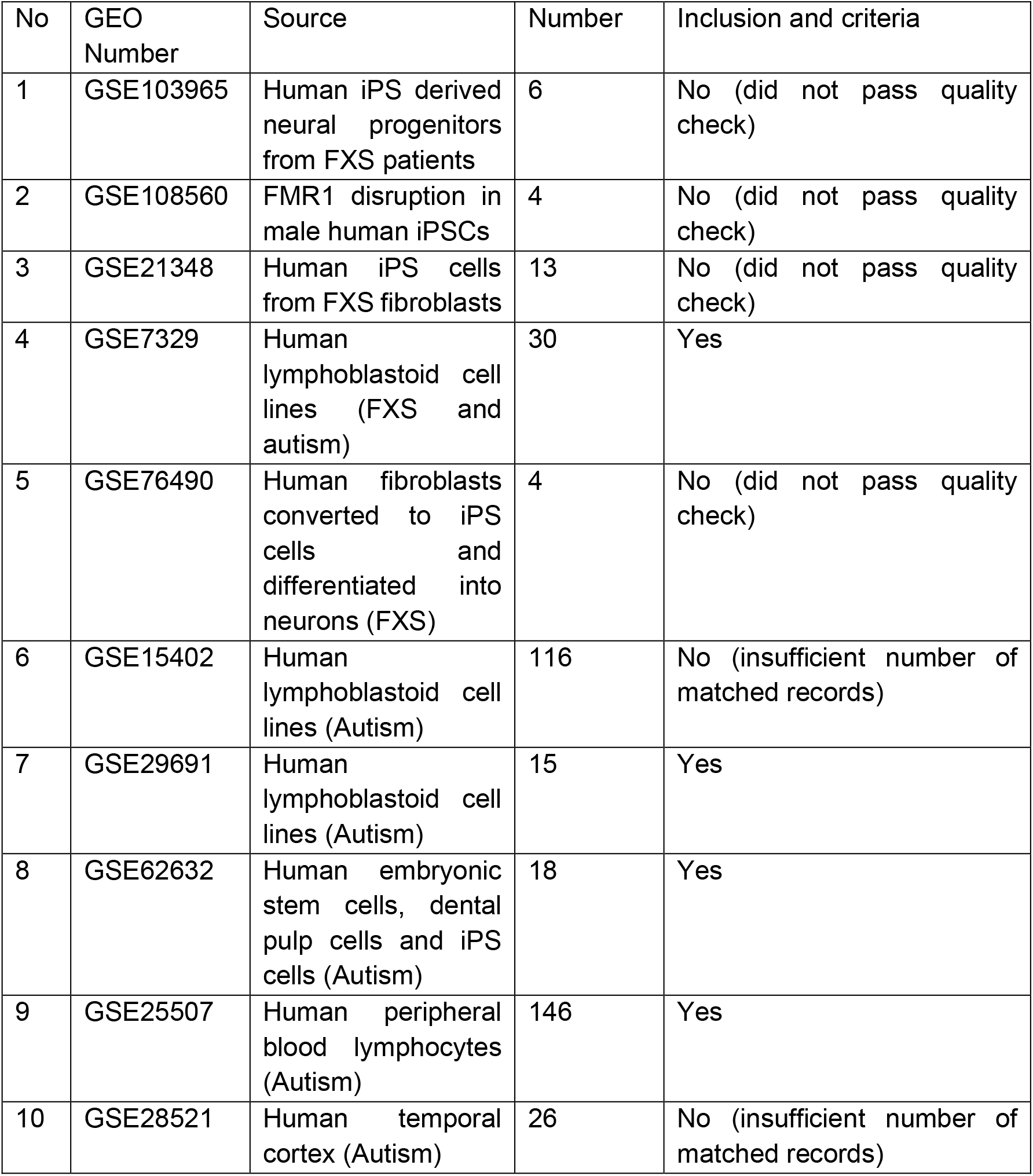
List of datasets (human) for FXS and ASD from NCBI GEO

### Data analysis to derive lists of differentially expressed genes (DEGs)

#### Individual analysis

The analysis of differentially expressed genes (DEG) for each dataset was performed with “Limma” package (linear models for Microarray data, a Bioconductor package), in R programming. Genes meeting the chosen criteria (P value) were selected as DEGs. The method involved the following steps: automated download of microarray datasets using getGEO to obtain expression values, background correction and normalization, fitting each probe set to a linear model with subsequent empirical Bayes adjustment. To identify DEGs between “disease” and “control”, Student’s t-test was performed. Fold change values were calculated, adjusted P-values were obtained. An adjusted P-value of P<0.01 was used as the criterion to identify DEGs.

#### Meta-Analysis

Chosen datasets were downloaded and the probe IDs were converted into gene symbols. For genes with multiple matched probe Ids, the average value of the probes was taken into account and finally converted to the format prescribed by MetaDE. All the datasets were read using MetaDE and merged using MetaDE.merge, to form a meta dataset containing the common gene profiles. For further analysis, using MetaDE.filter, 30% of un-expressed or un-informative genes were filtered out and the rest of the genes were used. Moderate t test was used to determine the difference in gene expression in each dataset and Fishers method was used to combine the P values. Benjamini-Hochberg method FDR, an adjusted P value <0.001 was used as the selection criteria of the DEGs.

### Random forest

Data pre-processing: To observe the performance of Random forest two types of data were used. 1. The raw values of gene expression data set 2. The normalized gene expression data were used, both having the gene names as features (columns) and the samples as the rows. Model: The random forest model was implemented by Python’s SKLEARN library. It creates multiple CART trees based on “bootstrapped” samples of data followed by voting approach of predictions leading to uncorrelated trees. After plotting OOB error estimate “100 trees” were chosen as n_estimators. Hence “n_estimators” here 100 bootstrap samples were drawn from the original data and for each bootstrapped data a tree was grown. At each node of the tree the “max_features (auto)” were randomly selected for splitting and the tree was grown until the terminal node had no lesser cases than “min_samples_leaf” (2). Then from all the trees the prediction was aggregated through voting approach. Subsequently OOB error rate was computed by using the data not present in the bootstrap samples. To reduce the classification error and to provide the best branch split of the trees gini index was used as the impurity measure metric.

The statistical formula of gini index is:

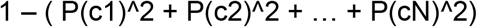

where p stands for probabilty and c stands for classes. Cross validation on our data was skipped since random forest has inbuilt bootstrap sampling and for hyper parameter tuning “Grid search CV” was used to get the best hyper prameters.

To validate the stability of the genes which were picked by variable importance and to understand the redundancy of the genes we performed bootstrap sampling, built 5-7 models and the variable importance (gene list) of bootstrap models were downloaded for a comparison study, shown in Fig. S3. The models were built with bootstrap sampling and different n_estimators, and all of the genes with variable importance greater than zero were chosen (to have a wide scope and large number of genes). The overlaps in these gene lists between models were analysed (Fig S3B) to assess model stability. The final gene list was found to be preserved in all other models (83% gene overlap) thus confirming good performance.

#### Hyper parameter tuning

was carried out in two steps here. The first parameter the “number of estimators” was selected through the OOB (out-of-bag) error estimate (Fig S1). From the OOB error estimate, we found that at and beyond the n_estimator value of 100, the error rate was reduced and maintained. Therefore, considering the computational cost, a value of 100 was chosen as the n_estimator. “GRIDSEARCH CV” was used as a second criterion to find the other hyper parameters And the best parameters were selected and used for the final model building. The final set of hyperparameters were as follows:

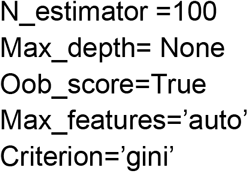

#### Support vector machines (SVMs)

To implement SVM, we chose to use sklearn feature selection “SelectFromModel” a meta transformer which selects features (genes) based on importance weights. SVM learns to differentiate between control and disease class by learning from examples, making it a pattern recognition algorithm. The SVM classifier was trained by 80% of data for our problem, that is these data points where taken to a high dimensional space and we tried finding the best hyperplane to classify. To do this the Kernel function was selected through GridSearch CV and “linear” kernel [k(x1,x2)=x1.x2] was selected as the best option as expected, based on nature of our data which is linearly seperable. The hyperparameter “c” was tuned using various options and 0.1 came out as best option for this problem. The results of the confusion matrix and classification reports, after tuning, are shown in Figure S4.

#### PCA

Principal component analysis (PCA) is a variance maximizing task, therefore standardization of gene expression values was carried out. The standardized data consists of only the features and not the dependent variable or outcome. For this study, PCA was implemented through the Python SKLEARN library. The number of principal components were given as the total components shaped as described “n_pcs = model.components_.shape[0]” leading to 209 components discovered as the direction of maximal variances in the data.

## Results and Discussion

### Identification of differentially expressed genes (DEGs) in FXS/Autism

In order to identify a gene signature that captures the changes that are strongly correlated with FXS, we started with transcriptomics or microarray data acquired from FXS patients. From NCBI GEO, we identified five FXS datasets, of which only one had sufficient sample number and data quality to be included in the analysis. FXS is part of the Autism spectrum, and a lot of the neurological symptoms in FXS patients match those seen in ASD patients. Therefore, on the assumption that that a core set of gene expression changes may be at the heart of both these syndromes, we expanded our scope to include autism datasets, of which we found three additional ones which were of sufficient quality and match between datasets, to be included in the current analysis.

Our goal was to use the gene expression information from these datasets, to compute a robust signature of “differentially expressed genes” (disease DEG) which would be characteristic of the core symptoms of FXS and autism (Figure 1). Several computational methods were used to determine this signature.

1. **Individual analysis**: Each dataset was analysed individually as described in the methods, and a list of genes with significant changes in expression between the control and disease samples, was found. At this stage, if the number of gene features present in the dataset was low, or if the magnitude of gene expression changes observed was not satisfactory, the dataset was not included in the combined analysis. Individual analysis was used to compute the fold-change in gene expression between disease and control samples.
2. **Meta Analysis:** Meta-analysis of gene expression data was done using the METADE package in R (MetaDE: Microarray meta-analysis for differentially expressed gene detection) (18), which implements 12 major meta-analysis methods for differential expression analysis. The output was in the form of a list of gene names, and an associated statistical significance measure (p-value) with meta.stat, meta.pvalue and meta.fdr and this list was considered the MetaDE-DEG signature.
3. **Gene selection using Machine Learning (ML) algorithms**: Machine learning can be used for gene profiling and selection. Feature selection, feature elimination and extraction algorithms were used to achieve the “gene selection” objective for this study. Feature selection algorithms are separated into three categories: i) Filters, which extract features from the data without any learning involved; ii) Wrappers, that use learning techniques to evaluate which features are useful and iii) Embedded techniques which combine the feature selection step and the classifier construction. Feature extraction creates new variables as combinations of others to reduce the dimensionality of the selected features. Hence, gene selection can also be considered a process of dimension reduction which allows one to narrow down the list of “important” genes. In this study, three different algorithms were used to find the disease DEGs. These were Principal Component analysis (PCA), Random forest (RF) and support vector machine (SVM). We chose these algorithms because of their use and proven track record with biological data(19–23). We selected three algorithms, each working on a different basis: RF-a tree based ensemble algorithm, PCA-a dimensionality reduction algorithm and SVM-a supervised algorithm taking the data into a higher dimension. The purpose of this pipeline was to understand which algorithm performs the best for these specific datasets and has the best classification scores.
  **3a. Random forest** is an ensemble algorithm for classification and regression developed by Leo Breiman, which can also be used to determine a list of “important features” along with “variable importance”. The datasets listed in Table 1 were used, and the results on the evaluation set of the final model after hyper parameter tuning are shown in Figure S2A (F1 score **2*((precision*recall)/(precision+recall)))**) and the area under the curve (AUC) in Figure S2B. These metrics were used to assess performace, and the model appeared to have performed well with AUC = 93% and F1 score 90%. Since the intended objective in this problem was to find the significant genes (important features) we considered this as our best performing model and the significant genes were extracted from this model using inbuilt feature called “variable importance”.
  **3b. Support vector machine (SVM)** is a supervised algorithm used for both classification and regression, but is preferred for classification due to its powerful classifiers built through kernel function and hyperlane concept. It is an ideal algorithm for high dimesional data such as gene expression values for a large set of genes/features. F1 score (**2*((precision*recall)/(precision+recall)))** and area under curve (AUC) were used as metrics to check the model performance, the model appeared to have performed good with AUC = 76% and F1 score 76%. The list of genes contributing to disease classification were extracted from the meta estimator “select from model “ with threshold mean and the details of significant genes are presented in Figure 2.
  **3a. PCA:** Principal component analysis is a feature extraction algorithm and unsupervised learning approach, which projects the original data into a low dimensional feature space and constructs the new dimensions. PCA is traditionally used in high dimensional data for reducing computational expense, increased error rate and noise and can also be used for gene selection. The high dimensional gene expression values can be reduced to low dimensional feature space with minimal loss of information, using this simple yet powerful unsupervised algorithm. PCA is one of the frequently used methods for feature extraction of microarray data. In this study, using the aforementioned datasets, we obtained an output of 209 Principal components resulting in gene list of 209 genes or features. The first four principal components (out of 209) of the PCA model are shown in Figure S5 and display the variance ratios captured.

**Figure 1:**
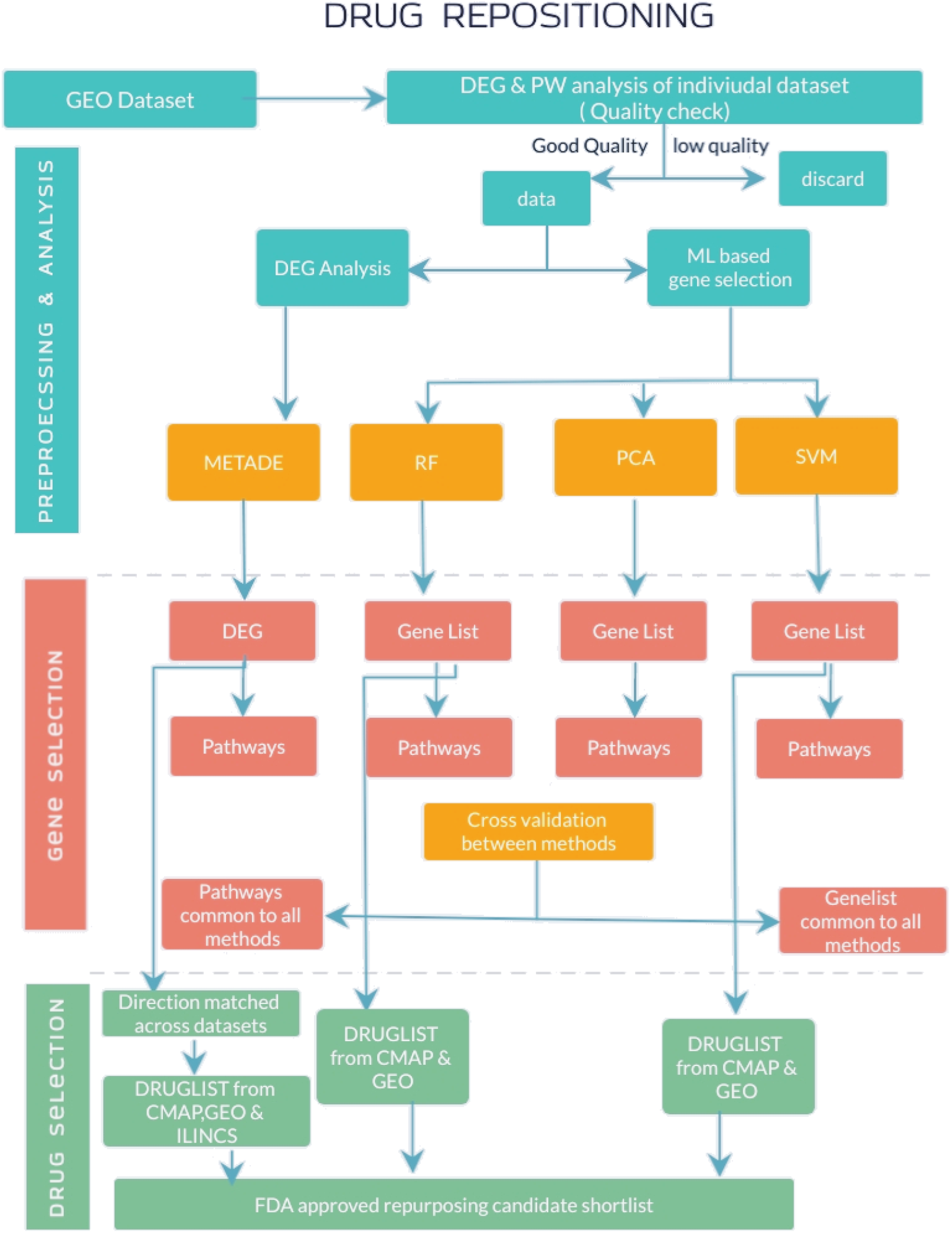
Overall Workflow

**Figure 2:**
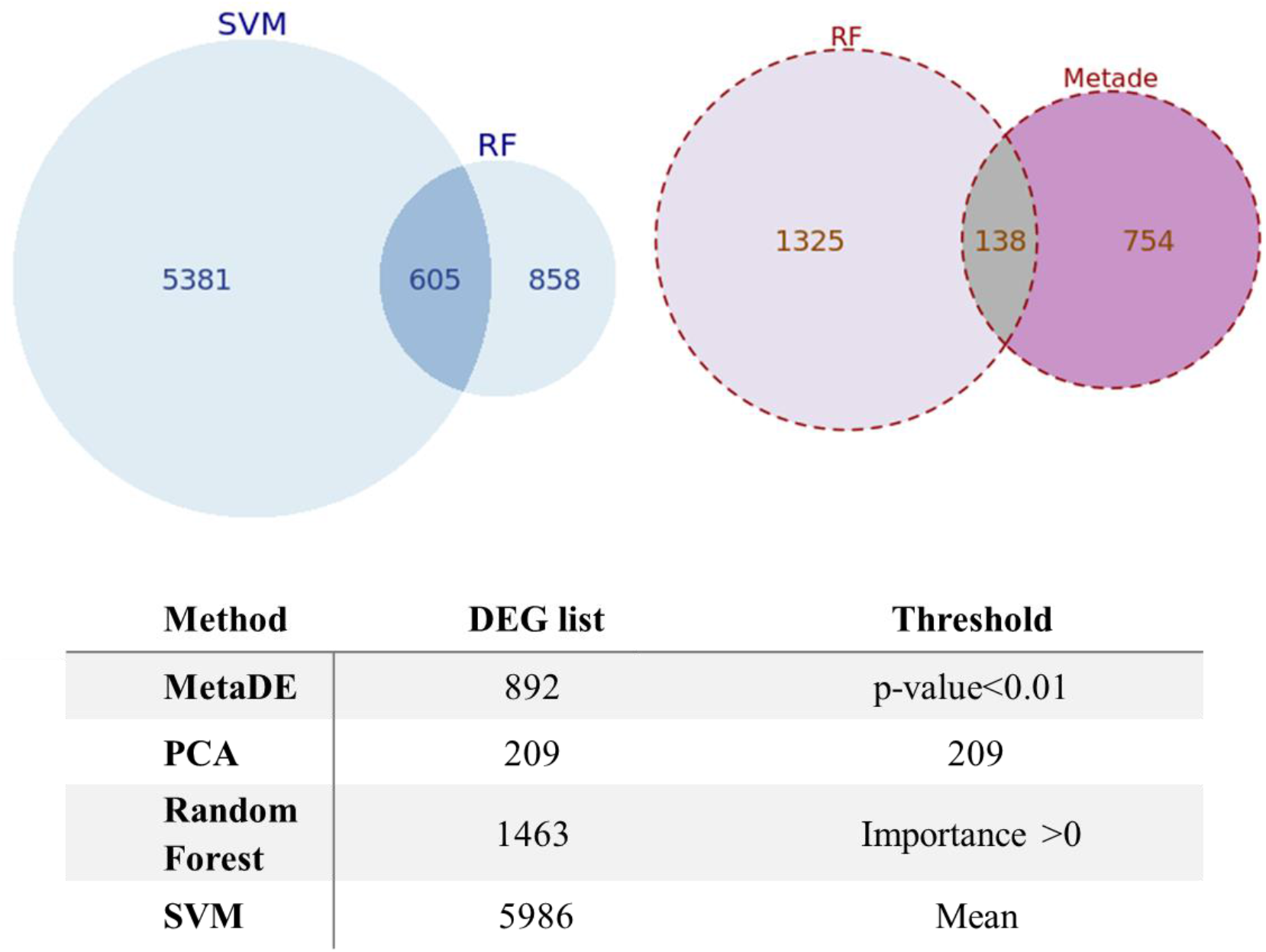
Summary of DEGs for FXS/ASD obtained using various computational methods

Among the machine learning algorithms, the performance of the RF algorithm was better than the other two in terms of classification, which implies that random forest and the “feature importance” assigned by the algorithm is a good metric to distinguish “control” and “disease” data. In general, ensemble algorithms are known for better performance and the same was true in the current study.

The number of genes that comprised the disease signature as determined by each of the methods described above, and the extent of overlap between them, is summarised in Figure 2. The DEG lists identified by Meta-analysis and machine learning, did not overlap significantly since the rationale behind each of the methods is different. However, since we had no way of knowing which method is better, we used these each of these gene lists to identify differentially affected pathways as the next step.

### Enriched pathways in FXS/Autism

Pathways represent groups of genes which are functionally linked, and together contribute to a cellular process or activity. A change in the expression of individual gene in a pathway may affect the function of the entire pathway, and conversely, dysregulated pathways may be manipulated by targeting any or several of the individual genes in a pathway. Therefore we sought to find the pathways that map to the DEG sets, which may characterize the FXS/autism disease state. The Enrichr tool (24) was used to identify enriched pathways in each of the identified DEG lists, using the KEGG 2019 database as a source of pathway information. The number of pathways identified for each gene list is indicated in Figure 3. We find that despite the poor overlap among the gene lists identified by MetaDE and RF, a majority of the enriched pathways are common. From this, we understand that a) no matter which algorithm is used, the core functional alteration present in the disease is captured by way of dysregulated pathways and b) using DEG lists derived from a single method as a basis for studying disease biology or therapeutic strategies may lead to incomplete or variable conclusions.

**Figure 3:**
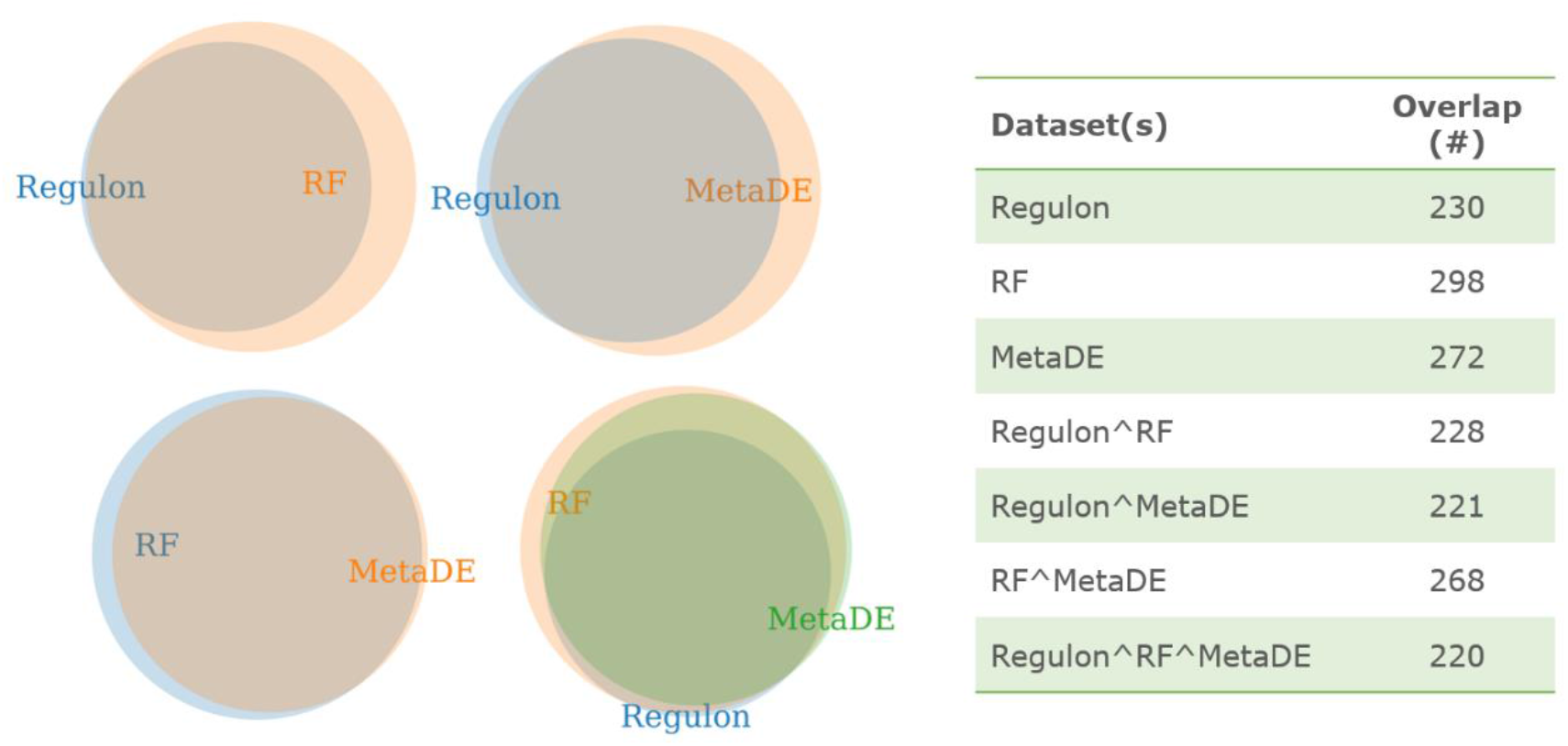
Venn diagrams to illustrate the overlap between pathways enriched in the gene lists from MetaDE, RF and regulons.

### Linking FMRP function to DEGs

Darnell et al reported the association of over 800 mRNAs with FMRP in neurons (5,25). As a protein known to interact with the ribosome and play a role in regulating translation elongation, it is likely that FMRP’s interaction with these mRNAs (referred to hereon in as FMRP-regulons) is probably to control their translation. FMRP mediated translation regulation has been demonstrated only for some of these target mRNAs, however it seems likely that several more could be regulated by FMRP in a similar manner. Therefore we looked for pathways enriched in these FMRP-regulons, and found that a significant number overlapped with the FXS pathways identified on the basis of DEGs (Figure 3). To summarise, the DEGs identified on the basis of the gene expression profile of human FXS and autism patients, and the targets of translation regulation by FMRP as identified in neurons, appear to be connected via the same pathways. A list of the top 20 enriched pathways are shown in Table 2. These results strengthen our inference that the DEG analysis has indeed identified a core set of features (genes and pathways) which are central to the nature of this disease, and may therefore be the most logical starting point for therapeutic intervention.

**Table 2:** Enriched pathways identified on the basis of disease DEGs

| **Enriched pathway** | **Overlap**<br>**(Regulon)** | **Overlap**<br>**(DEG)** |
| --- | --- | --- |
| *Synaptic vesicle cycle* | 22/78 | 14/78 |
| *Adrenergic signalling in cardiomyocytes* | 26/145 | 19/145 |
| *Endocytosis* | 24/244 | 28/244 |
| *Dilated cardiomyopathy (DCM)* | - | 13/91 |
| *Dopaminergic synapse* | 26/131 | 17/131 |
| Fc gamma R-mediated phagocytosis |  | 17/91 |
| Human cytomegalovirus infection | 24/225 | 26/225 |
| Endocytosis | 24/244 | 25/244 |
| Human papillomavirus infection | 25/330 | 30/330 |
| Pathways in cancer | 35/530 | 42/530 |
| Cholinergic synapse | 18/112 | 14/112 |
| Hepatocellular carcinoma | 17/168 | 18/168 |
| MAPK signaling pathway | 27/295 | 26/295 |
| Sphingolipid signaling pathway | 11/119 | 14/119 |
| Parathyroid hormone synthesis, secretion and action | 15/106 | 13/106 |
| Estrogen signaling pathway | 16/137 | 15/137 |
| Cushing syndrome | 16/155 | 16/155 |
| Long-term potentiation | 18/67 | - |
| Autophagy | 18/128 | 13/128 |
| HIF-1 signaling pathway | 11/100 | 11/100 |
| Dopaminergic synapse | 26/131 | 13/131 |
| cGMP-PKG signaling pathway | 23/166 | 15/166 |
| Focal adhesion | 16/199 | 17/199 |
| Amphetamine addiction | 13/68 | - |
| Wnt signaling pathway | 16/158 | 14/158 |

### Drug selection based on the disease DEG and enriched pathways

Just as a genetic change that leads to a disease can be correlated to a gene expression signature, drug treatment is also expected to result in predictably associated gene expression changes. The Connectivity map (CMAP) and Library of Integrated network-based signatures (LINCS) databases provide a readily accessible catalogue of such drug induced gene expression profiles, based on a panel of cell lines, for a large number of small molecules and other perturbagens. Similarly, the NCBI GEO database also provides access to gene expression data from drug treated samples (cell lines, animal models, human samples). These databases can be queried using tools such as Gene2drug (CMAP)(26), iLINCS (LINCS) (27) and Enrichr (GEO), wherein the DEG list is used as the input to extract drugs which result in similar or reverse DEG profiles. In this study, the MetaDE DEG and RF DEG lists were used to query all three databases, the resultant list of small molecules were filtered on the basis of current FDA approval and are presented in Table 3. The MetaDE DEG list is associated with a direction of change in gene expression, and was therefore used to identify only those treatments which reversed the change. Since the RF DEG list is not associated with direction (due to the nature of the algorithm), drugs that were associated with the DEG irrespective of direction, were identified. The master list could therefore include compounds which will ameliorate or worsen the phenotypes associated with FXS/ASD.

**Table 3:**
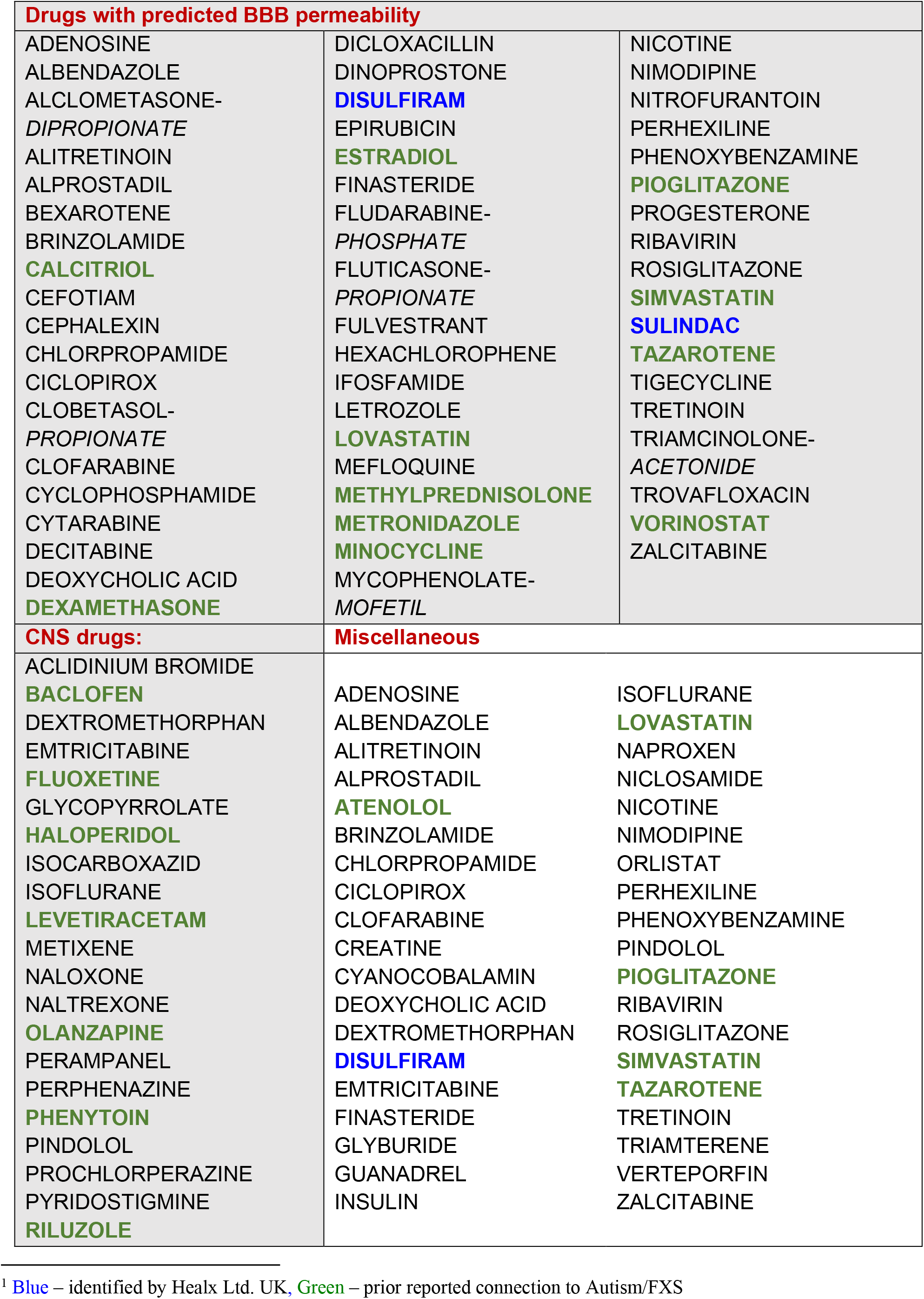
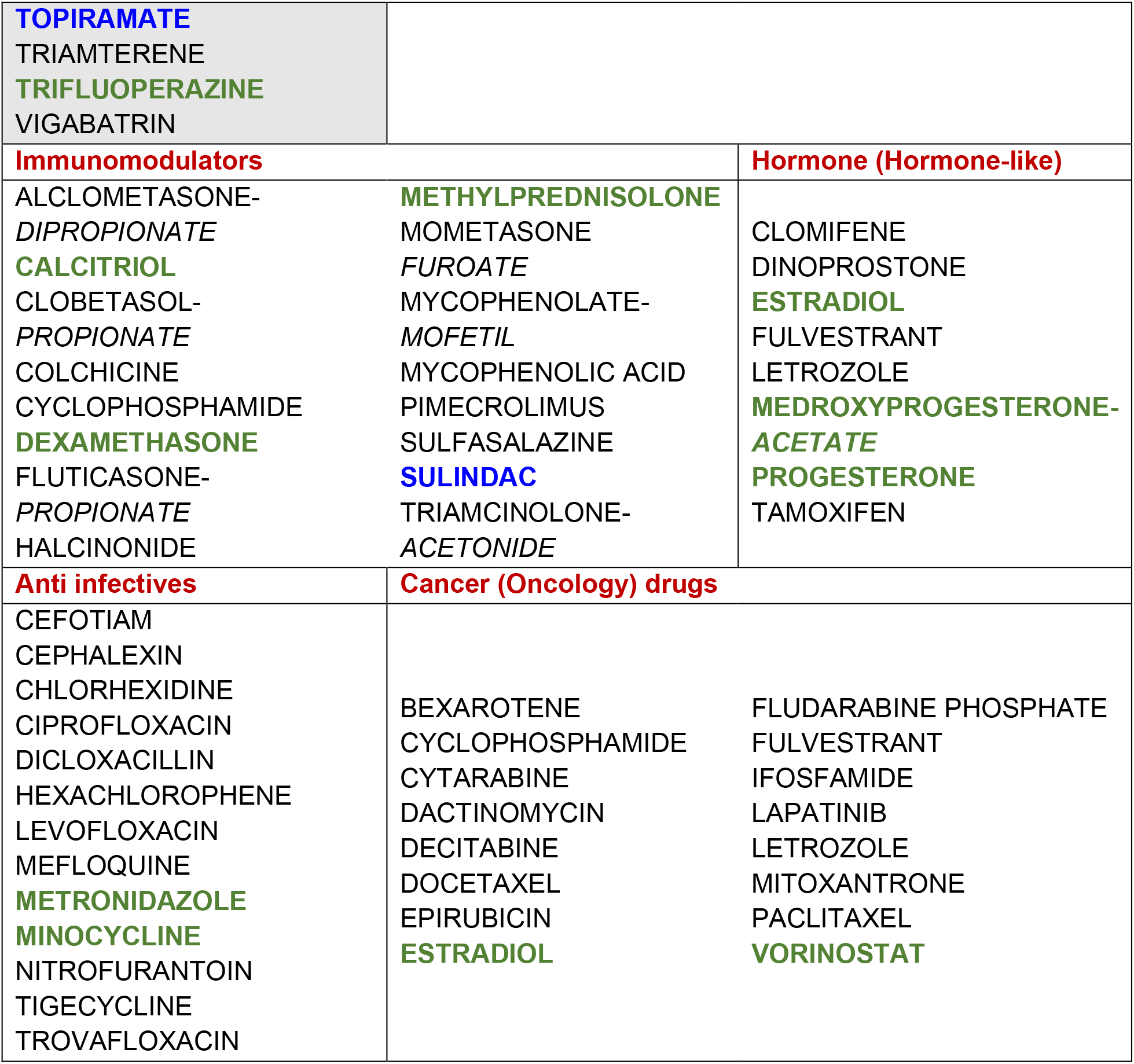
List of FDA approved drugs identified by the pipeline (organized by therapeutic class)^1^:

### Characteristics of identified drugs

Among the drugs identified on the basis of multiple DEG lists, over 30% are known to cross the blood brain barrier or are approved for a neurology indication; over 25 drugs have a reported previous association with FXS and/or ASD in the published literature; at least 3 have been previously identified in an AI-based repurposing study by Healx Ltd. UK, and reported to improve outcomes in the FXS mouse model (Table 3). Therefore there is a high likelihood that one of these drugs could potentially be repurposed to treat ASD symptoms. A mechanistic link to FXS/autism is apparent for some drugs and a few examples are as follows: Chlorpropamide is an oral antihyperglycemic agent which is known to increase the secretion of vasopressin, which may have a positive impact on social functioning (28); Tazarotene is a retinoid compound approved for skin disorders, and through its action on the RAR family, could influence autistic phenotypes (29); Pioglitazone, a PPARγ agonist has just recently been shown to improve behavioural deficits in FXS mice by limiting the excessive diacylglycerol signalling in the brain (30) and low dose Trifluoperazine treatment, acting via the PI3K/Akt pathways has been proposed as a potential therapeutic for FXS based on in silico analysis of gene expression data and studies in mice (31). The rationale for drugs belonging to certain drug classes such as immunomodulators and anti-infectives, can be explained on the basis of the link between autism, neuroimmune dysregulation and microbial dysbiosis (32). Based on these examples and reports from the literature, it appears that the candidates picked by the DREAM-RD pipeline have a high likelihood of positive impact for FXS/autism therapy.

While these small molecules were identified on the basis of the changes they induce at the level of gene expression, practical considerations such as blackbox warnings, acute toxicities and side effects may preclude their use for FXS treatment. A filtered list without these incompatible drugs is shown in Table 4, and this could be the starting point for testing molecules for efficacy in FXS models.

**Table 4:** Shortlist of drugs identified by the pipeline without severe blackbox warnings or acute toxicities.:

| **Drugs with no blackbox warnings or chronic toxicity** |  |
| --- | --- |
| ACLIDINIUM BROMIDE<br>ALBENDAZOLE<br>ALCLOMETASONE-DIPROPIONATE<br>ALITRETINOIN<br>BACLOFEN<br>BRINZOLAMIDE<br>CALCITRIOL<br>CEPHALEXIN<br>CHLORPROPAMIDE<br>CICLOPIROX<br>CLOBETASOL-PROPIONATE<br>CLOMIFENE<br>COLCHICINE | HEXACHLOROPHENE<br>INSULIN<br>LETROZOLE<br>LEVETIRACETAM<br>LOVASTATIN<br>METHYLPREDNISOLONE<br>METIXENE<br>MINOCYCLINE<br>MOMETASONE FUROATE<br>NALOXONE<br>NALTREXONE<br>NICLOSAMIDE<br>NICOTINE |
| CREATINE | NITROFURANTOIN |
| CYANOCOBALAMIN | ORLISTAT |
| DECITABINE | PERHEXILINE |
| DEOXYCHOLIC ACID | PINDOLOL |
| DEXAMETHASONE | PYRIDOSTIGMINE |
| DEXTROMETHORPHAN | RILUZOLE |
| DICLOXACILLIN | SIMVASTATIN |
| DINOPROSTONE | SULFASALAZINE |
| FINASTERIDE | TAZAROTENE |
| FLUTICASONE- PROPIONATE | TOPIRAMATE |
| FULVESTRANT | TRIAMCINOLONE- ACETONIDE |
| GLYBURIDE | VERTEPORFIN |
| GLYCOPYRROLATE | VORINOSTAT |
| GUANADREL | ZALCITABINE |
| HALCINONIDE |  |

In this study, we have identified both a list of characteristic gene expression level changes that represent the altered cellular state of FXS/Autism, and a list of accompanying small molecules that could perturb this state and potentially restore the imbalance. The next steps will involve conducting screens in appropriate models to test the efficacy of the drug in reversing symptoms, and validation of the underlying mechanism. While there are many subsequent steps required for validation and drug development before a clinical outcome may be reached, such a strategy is likely to be more successful than both the completely random and expert-intuition based approaches which have been in place thus far since it allows for the inclusion of the non-obvious, yet significant changes found across multiple diverse datasets.

Our approach, once validated, can easily be applied to any number of diseases, rare or otherwise. One major limitation is the availability of good quality data in sufficient quantity, from human patients. This deficit needs to be corrected and is possible with the rapidly reducing costs and increasing availability of transcriptomics facilities. The second caveat is the use of drug DEG profiles obtained from cell line experiments, which could be misleading and inappropriate especially for repurposing, where the effect on the whole organism is likely to contribute to the overall outcome. Therefore, generation of such drug induced profiles in model organisms, in a temporal as well as tissue specific manner is likely to add immense value and contribute to a greater chance of success during the transition to the clinic.

## Acknowledgments

We are very grateful to Sridivya Pantula for contributing to the collection and classification of drug information, and to Dr.Kiranam Chatti, Dr. Kishore Parsa and Dr. Srinivas Oruganti for discussions on methodology and advice and comments on the manuscript.

## Competing interests

The authors declare no competing interests.

## Funding

This study was supported funding from Dr.Reddy’s Institute of Life Sciences, Hyderabad.

## Author contributions statement

K.A and A.S. conceptualized the study and designed the pipeline. K.A. was responsible for exploratory data analysis including data collection and cleaning, programming, algorithm selection and implementation. A.S was responsible for data selection, curation and guidance during pipeline implementation. K.A and A.S analyzed the results and wrote the paper.

## Supplementary information

**Figure S1:**
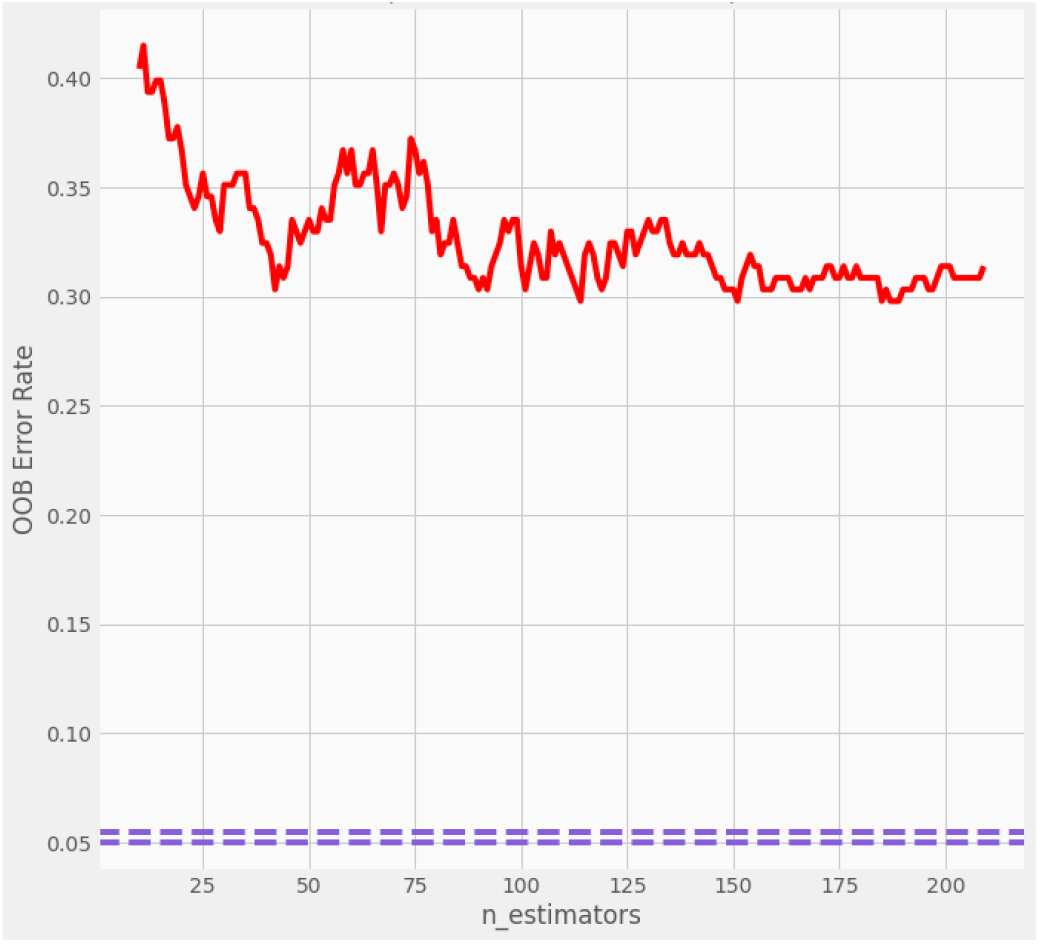
OOB error rate across various forest sizes (10-200)

**Figure S2:**
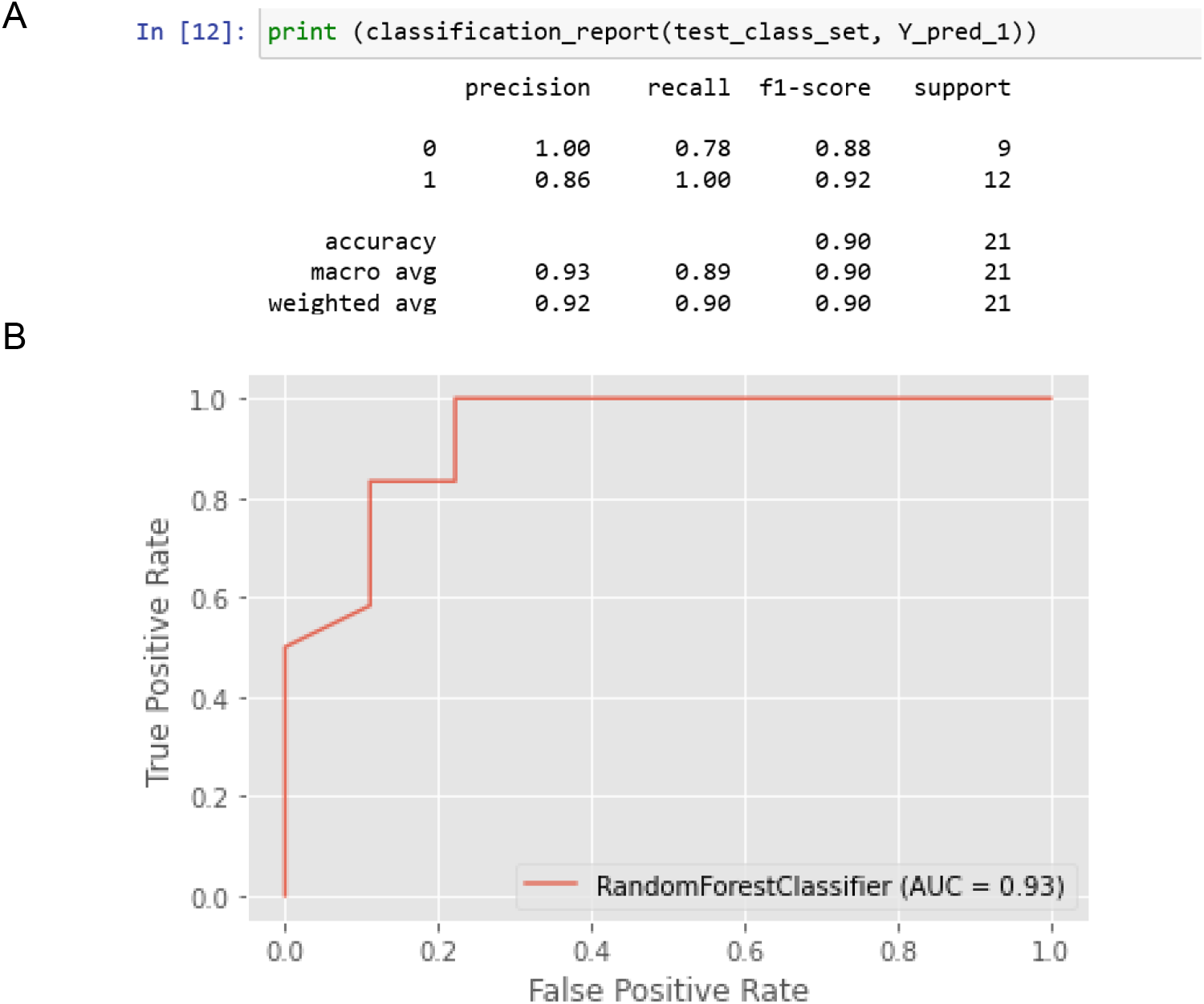
Evaluation set for the final RF model

**Figure S3:**
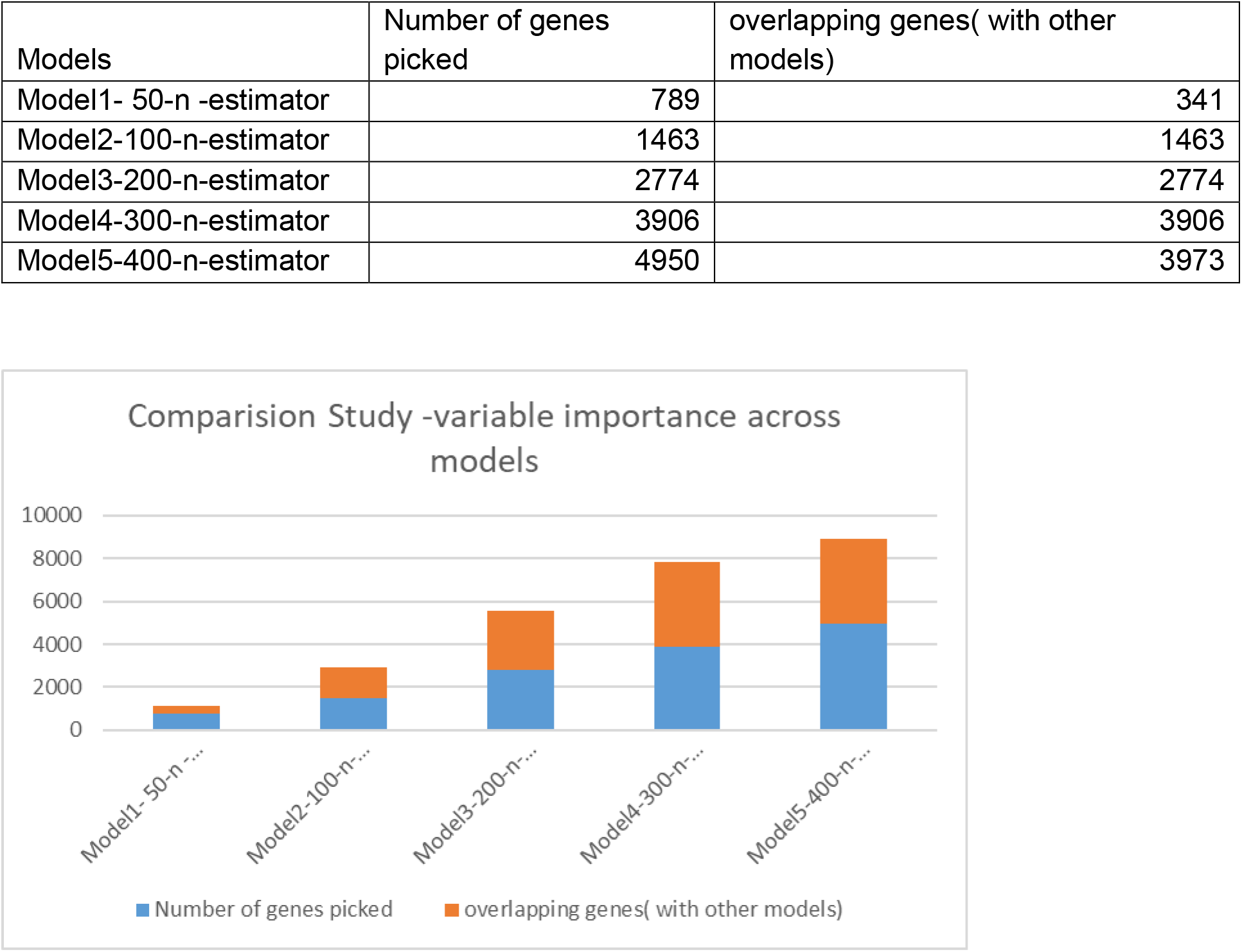

**Figure S4:**
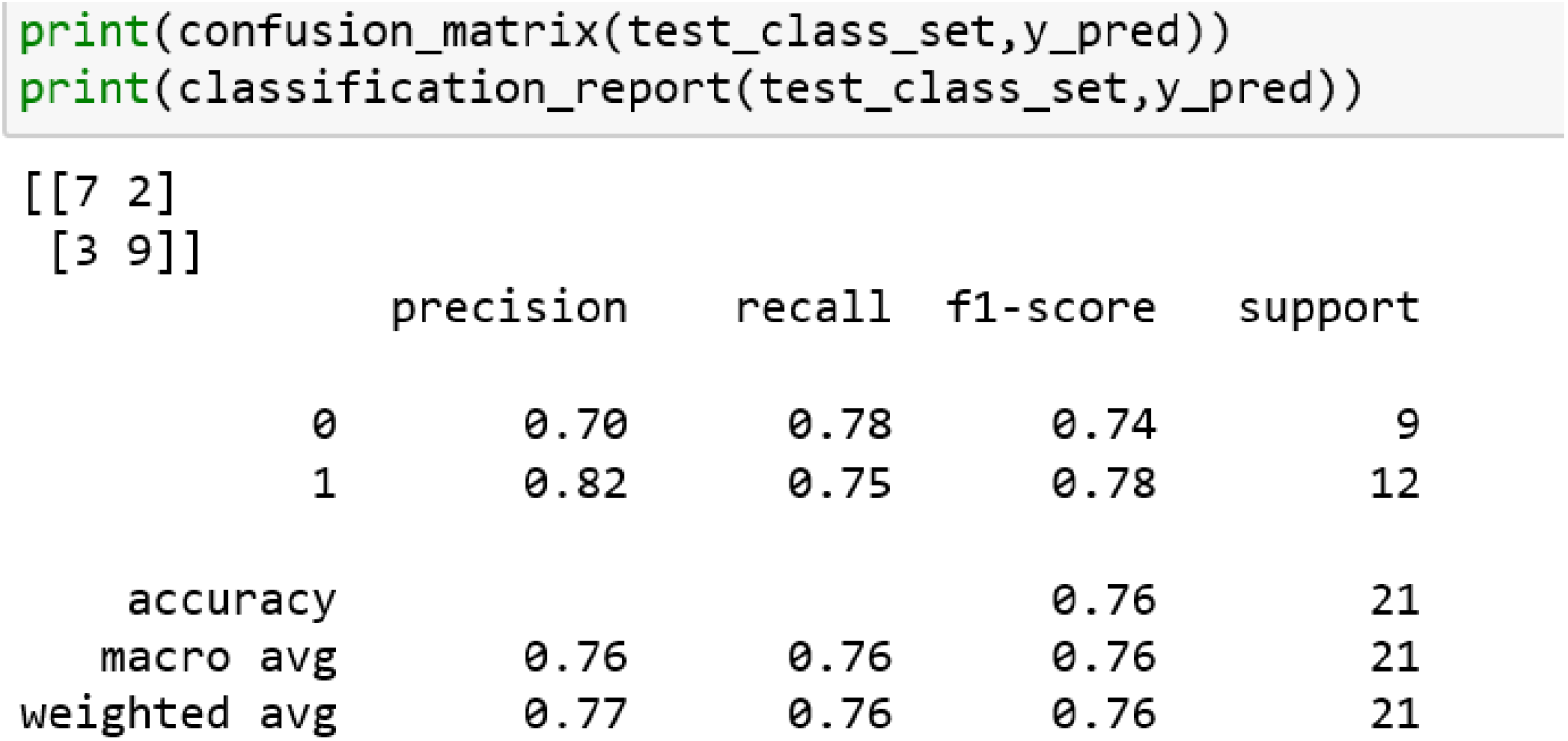
Evaluation report for the SVM algorithm.

**Figure S5:**
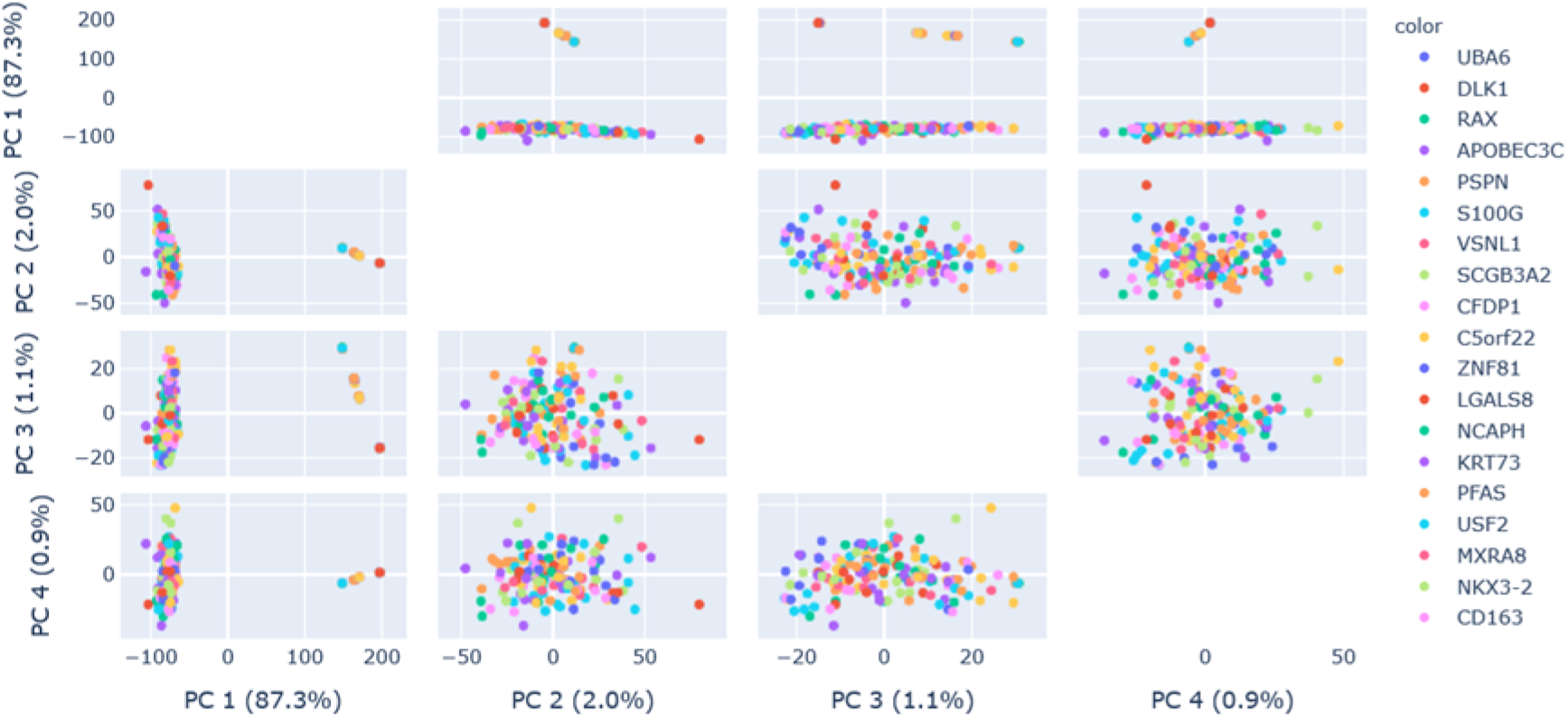
Variance ratios in the first four PCs

## Notes

### Competing Interest Statement

The authors have declared no competing interest.

